# SDF-1/CXCR4 inhibition prevents paradoxical generation of cisplatin-induced pro-metastatic niches

**DOI:** 10.1101/2020.10.26.355057

**Authors:** Giulia Bertolini, Valeria Cancila, Massimo Milione, Giuseppe Lo Russo, Orazio Fortunato, Nadia Zaffaroni, Monica Tortoreto, Giovanni Centonze, Claudia Chiodoni, Federica Facchinetti, Francesca Giovinazzo, Massimo Moro, Chiara Camisaschi, Alessandro De Toma, Crescenzo D’Alterio, Ugo Pastorino, Claudio Tripodo, Stefania Scala, Gabriella Sozzi, Luca Roz

## Abstract

Platinum-based chemotherapy remains widely used in advanced non-small cell lung cancer (NSCLC) despite its ineffectiveness in long-term control of metastasis.

Here, we uncover the interconnected pathways subtending cisplatin-induced metastasis promotion.

We report that cisplatin treatment of tumor-free mice results in bone-marrow expansion of CCR2+CXCR4+Ly6C^high^ inflammatory monocytes (IM) concomitantly with increased levels in the lungs of stromal SDF-1, the CXCR4 ligand. In experimental metastasis assays, cisplatin-induced IM favor tumor cells extravasation and expansion of CD133+CXCR4+ metastasis initiating cells (MICs), facilitating lung metastasis formation. At the primary tumor, cisplatin reduces tumor size but induces tumor release of SDF-1 triggering MICs expansion and recruitment of pro-invasive CXCR4+ macrophages. Co-recruitment of MICs and CCR2+CXCR4+ IM at SDF-1-enriched distant sites also promotes spontaneous metastasis. Combination treatment with a CXCR4 inhibitor prevents cisplatin-induced IM/MICs recruitment and interaction thus precluding metastasis overgrowth. Finally, we observe in NSCLC patients’ specimens that SDF-1 levels are higher in platinum-treated samples and correlate with worse outcome.

Our findings suggest a possible novel combination therapy based on CXCR4 blockade to control metastatic disease, paradoxically promoted by cisplatin.

## Background

Lung cancer represents the first cause of cancer-related mortality worldwide [1]. Platinum-based chemotherapy represents the standard of care for neo-adjuvant/adjuvant treatment of advanced stage NSCLC patients, alone or in combination with immunotherapy [2–6]. Nevertheless, long term effectiveness of standard chemotherapy is mined by potential detrimental effects such as promotion of tumor relapse and metastasis spread [7,8].

In preclinical settings, chemotherapy treatments, despite showing effective control of primary tumors, may paradoxically stimulate pro-metastatic effects mediated by host inflammatory responses to cytotoxic damages [9,10]. Chemokines released by damaged stromal/tumor cells after chemotherapy can promote tumor cell dissemination, survival and growth at distant sites [7,11,12]. However, how platinum compounds can orchestrate multiple pro-metastatic activities are incompletely understood.

The myeloid subset of C-C motif chemokine receptor type 2 (CCR2)^high^ inflammatory monocytes (IM) promotes early phases of metastasis development by fostering tumor cells extravasation at distant sites [13,14]. Chemotherapy can exacerbate CCR2+ IM recruitment and consequent metastasis formation by inducing tumor/stromal release of the ligand CCL2 [15,16]. Moreover, chemotherapy can induce the release of other chemokines such as SDF-1, the ligand for chemokine receptor CXCR4, resulting in recruitment of a subset of tumor associated macrophages (CD206+TIE2^High^ CXCR4^high^) with pro-angiogenic activity [17,18], favoring primary tumor re-growth and dissemination [19].

CXCR4, a G-protein coupled receptor (GPCR) expressed on multiple cell types including lymphocytes, hematopoietic stem cells, endothelial cells, fibroblasts and cancer cells [20,21], is implied in several mechanisms favoring tumor progression and metastasis [20,22,23]. CXCR4 inhibitors are under investigation for treatment of refractory late-stage solid tumors with promising results [20,24,25].

We previously reported that in lung cancer the fraction of CD133+ cancer stem cells (CSCs) coexpressing the chemokine receptor CXCR4 can be spared by chemotherapy and possesses the highest ability to disseminate and initiate distant metastasis (metastasis initiating cells, MICs) [26–28]. Cisplatin treatment of lung cancer patient derived xenografts (PDXs), despite shrinking primary tumors, results in MICs enrichment and enhances metastasis formation that can be counteracted by CXCR4 inhibition [26].

Here we hypothesized a central role of the CXCR4/SDF-1 axis in orchestrating multiple chemotherapy-induced pro-metastasis effects in NSCLC and tested the effectiveness of a combination therapy with a novel peptide inhibitor of CXCR4 (peptide R), designed as an SDF-1 mimetic peptide [29,30], to counteract cisplatin promotion of prometastatic microenvironments and improve treatment efficacy.

## Results

### Cisplatin promotes expansion of bone marrow-derived inflammatory monocytes (IM) and their lung recruitment through SDF-1/CXCR4 activation

In light of pre-clinical evidence on the functional relevance of systemic pro-metastatic changes induced by chemotherapy, we initially investigated the direct effect of cisplatin in shaping the microenvironment and promoting the generation of a pre-metastatic niche in healthy tissues. We focused in particular on the CXCR4/SDF-1 axis which we previously identified as a crucial modulator of cancer stem cells dynamics in cisplatin-treated PDX models [26]. Tumor-free naïve SCID mice were treated with cisplatin alone or in combination with the CXCR4 inhibitor peptide R [29] and reactive changes were evaluated in the bonemarrow (BM) and lungs. After 72h, cisplatin caused the expansion of BM inflammatory monocytes (IM) (CD11b+Ly6C+Ly6G-), without substantial changes of other myeloid cells or NK cells (Fig. 1a, b). Notably, IM in cisplatin-treated group were 4-fold enriched in the subset co-expressing CXCR4 and CCR2. CXCR4 inhibition prevented IM expansion caused by cisplatin and the enrichment of the CXCR4+/CCR2+ subset (Fig. 1a, b). In the lungs, cisplatin-induced damages resulted in the release of several inflammatory cytokines including CCL2 and SDF-1, that was significantly impaired by CXCR4 inhibitor (Appendix Fig. S1a, b and Fig. 1c). Notably, SDF-1 levels at BM remained substantially unaffected by the treatment (Appendix Fig. S1c). Immunofluorescence analysis of lung tissues showed that SDF-1 production was predominantly contributed by interstitial PDGFRβ+ lung pericytes while only minimally associated to CD45+ elements, data confirmed by *in vitro* combination treatments of both human and murine endothelial cells (Fig. 1d and Appendix Fig. S1d).

**Figure 1.**
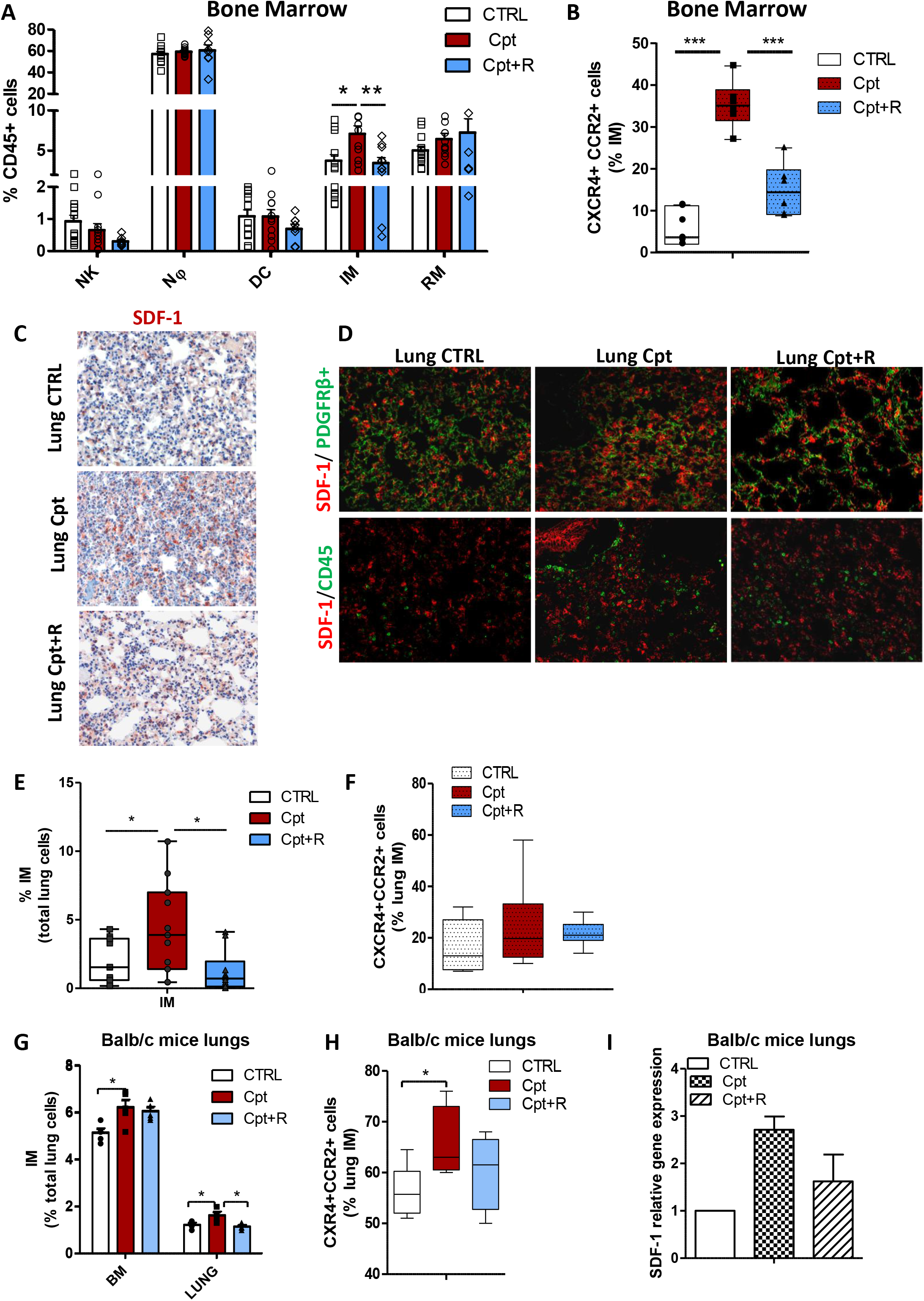
Cisplatin promotes lung recruitment of BM-inflammatory monocytes through SDF-1/CXCR4 activation. **A)** FACS analysis of DX5+ NKs and CD11b+ myeloid cell subsets in bone marrow of SCID mice 72h post treatment with cisplatin alone or in combination with peptide R (CXCR4 inhibitor). N=5 mice/group, two independent experiments. Bars are the mean value ± SE. *p≤ 0.05. Nφ= Neutrophils; DC=Dendritic Cells; IM=Inflammatory Monocytes; RM= Resident Monocytes. **B)** Frequency of CXCR4 and CCR2 positive inflammatory monocytes, detected in (A). ***p≤ 0.001. **C)** IHC for SDF-1 in lung tissue of SCID control and treated mice, 72h after administration of cisplatin alone and in combination with peptide R. **D)** Double immunofluorescence for SDF-1 and PDGFRβ+ stroma cells (on the top) or CD45+ immune cells (on the bottom) performed on lung tissue from the same experimental groups as in (C). **E)** Median percentage of inflammatory monocytes, gated total lung live cells, detected by FACS in lungs of mice 72h post treatment with cisplatin and combination therapy compared to untreated control. N=5 mice/group in two independent experiments. *p≤ 0.05. **F)** Frequency of CXCR4 and CCR2 positive inflammatory monocytes, detected in E. **G)** Percentage of inflammatory monocytes, relative to the gated live lung cells, evaluated by FACS in BM and lungs of BALB/c mice, 72h post treatments with cisplatin and combination therapy compared to untreated control. N=5mice/group. Bars are the mean value ± SD. *p≤ 0.05. **H)** Median frequency of CXCR4 +CCR2+ cells within gated inflammatory monocytes, detected in (G). *p≤ 0.05. **I)** Analysis for *sdf-1* gene expression performed in lung tissue of n=5 BALB/c mice treated with cisplatin or combination. Untreated mouse lung tissue was used as calibrator.

Consistently with the release of SDF-1 and CCL2, cisplatin increased the relative abundance of CCR2+ IM (2.3-fold change) expressing higher levels of CXCR4 then untreated controls; combination with the CXCR4 inhibitor prevented the increased of inflammatory monocyte to the lung (Fig. 1e, f).

Since inflammatory monocytes share similar phenotype with monocytic-myeloid derived suppressor cells (M-MDSC) [31], we also assessed the expression of a series of immunosuppressive (IL-10, Nos1, Nox2) and MDSC-related genes (CD49d and Arginase) but no modulation or even detection can be found in treated lung tissue (Appendix Fig. S1e), suggesting that the cisplatin-recruited CD11b+Ly6C+Ly6G-myeloid subset is likely represented by IM rather than M-MDSC.

Inflammatory status promoted by cisplatin and resulting increase of IM at BM and lung sites were mainly observed in acute response (72h after treatment) and alleviated during prolonged treatments (2-4 weeks) (Appendix Fig.1Sf-h).

Acute treatment (72h) with cisplatin of tumor-free immunocompetent mice confirmed the expansion of BM-derived IM and their recruitment to the lungs through a CXCR4-dependent mechanism (Fig. 1g-l). Cisplatin treatment did not significantly affect other myeloid or lymphoid immune cell compartments (Appendix Fig. S1i). Interestingly, no modulations in the expression of M-MDSC-associated genes or in T-reg frequency were observed in lungs after acute treatment with cisplatin (Appendix Fig. S1i, l).

Treatments of naïve SCID mice with other drugs commonly used for lung cancer treatment (Gemcitabine, Paclitaxel, Pemetrexed) failed to cause the expansion of BM derived IM (Appendix Fig. S2a) and to significantly increase stromal expression of SDF-1 and CCL2 cytokines in the lungs that abundantly chemoattracted CXCR4+CCR2+ IM (Appendix Fig. S2b-e).

Overall, these data indicate a specific role of CXCR4/SDF-1 axis in mediating cisplatin-induced inflammation.

### Inflammatory monocytes recruited by cisplatin support lung metastasis outgrowth

Next, to elucidate the impact of cisplatin-induced host inflammatory responses in metastasis promotion, the human metastatic lung cancer cell line H460 was inoculated in the tail vein of SCID mice 72h after administration of cisplatin (alone or in combination with peptide R). One week after injection, the number of H460 cells seeding the lungs was only slightly increased in cisplatin pre-treated animals but significantly enriched for MICs subset compared to untreated controls (Fig. 2a, b). Three weeks after injection, the cisplatin pre-treated group showed a massive increase in number and size of metastatic foci that also maintained MICs enrichment compared to untreated controls (Fig. 2c-e). Pre-treatment with peptide R prevented MICs survival and expansion caused by cisplatin (Fig. 2a-e).

**Figure 2.**
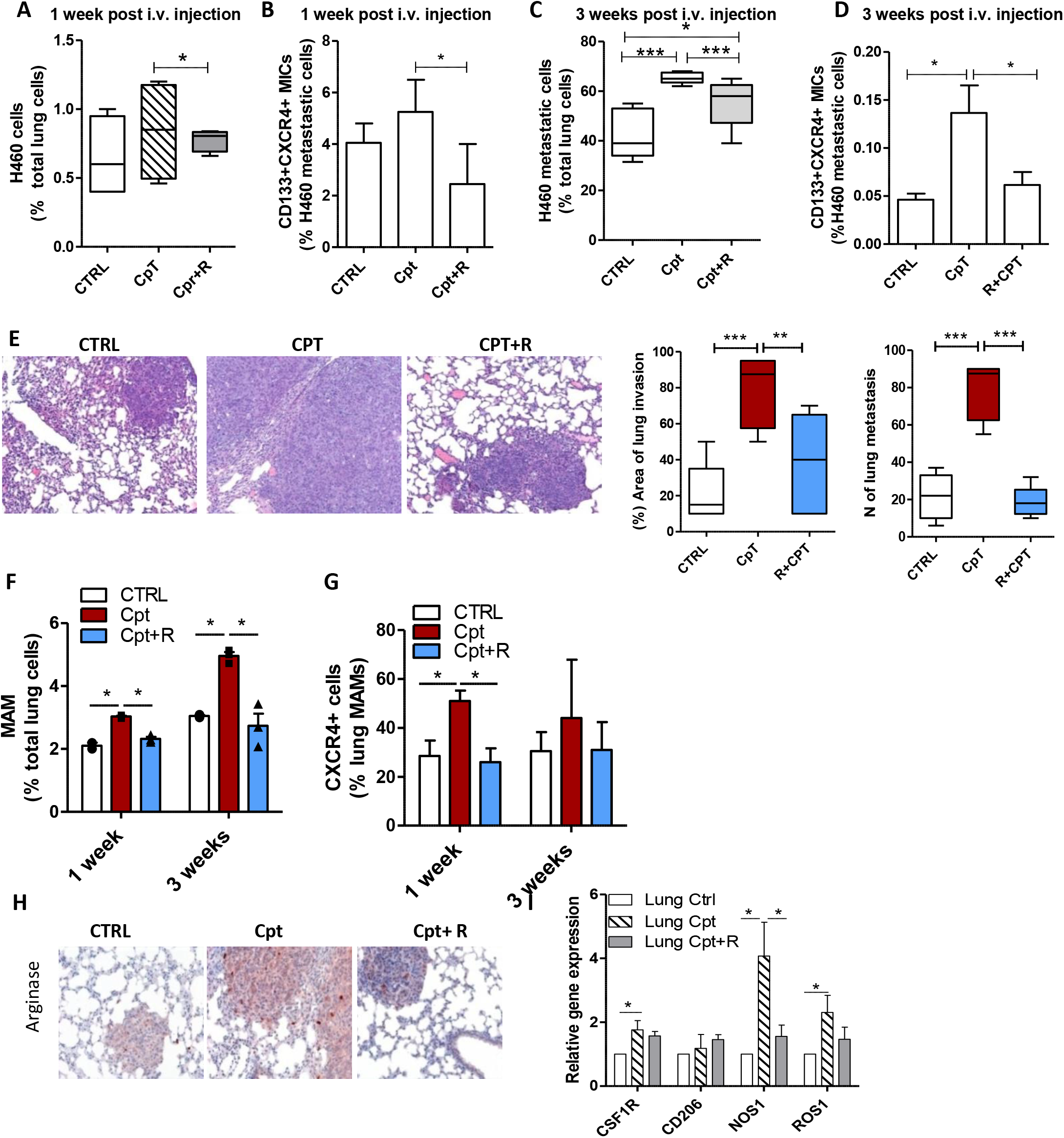
Cisplatin-activated microenvironment induces MICs selection and expansion via SDF-1/CXCR4 axis. **A)** Percentage of H460 cells surviving in lungs of SCID mice pre-treated with cisplatin or combination with CXCR4, 1 week after i.v. injection. Tumor cells were detected by FACS as live (7-AAD-) and murine HLAneg cells. N=3mice/group for two independent experiments. **B)** Frequency of CD133+CXCR4+ MICs within gated H460 cells in murine lung (as detected in A). N=3mice/group for two independent experiments. * p≤ 0.05. **C)** Median percentage of metastatic H460 cells detected by FACS in murine lung, 3 weeks after i.v. injection in SCID mice pre-treated with cisplatin or combination with CXCR4. N=6 lungs/group, for three independent experiments. * p≤ 0.05. ***≤ 0.001. **D)** Frequency of CD133+CXCR4+ MICs within H460 lung metastases, detected in (C). Bars are the mean value ± SD of N=6 lungs/group for three independent experiments. * p≤ 0.05. **E)** H&E of murine lungs, 3 weeks after H460 i.v. injection in SCID mice pre-treated with cisplatin or combination with CXCR4 (on the left) and histological quantification of the area of lung invasion and number of metastatic foci (on the right). N= 3mice/group, two independent experiments. ***p≤ 0.0001; **p≤ 0.001. **F)** Percentage of metastasis-associated macrophages (MAM) detected by FACS in murine lungs 72h post treatment with cisplatin and 1 and 3 weeks after H460 cells i.v. injection. Bars are the value ± SD. N=3mice/group, two independent experiments was performed for the 72h time point. * p≤ 0.05. **G)** Frequency of CXCR4+ cells in MAM detected in (F). Bars are the value± SD. N=3mice/group, two independent experiments was performed for 72h time point. *p≤ 0.05. **H)** IHC for Arginase+ MAM in lungs metastasis analyzed in (E), three weeks post treatment. **(I)** Real Time analysis of the same lung tissue for MAM-associated genes. Bars are the value relative to untreated control ± SD. N=3mice/group.

During lung metastasis development, we observed an increasing number of metastasis associated macrophages (MAM) (CD11b+CD11c-F480+GR1-)[32], highly expressing CXCR4, that remain more abundant in lungs of cisplatin pre-treated mice (Fig. 2f, g).

Accordingly, in metastatic lungs of cisplatin pre-treated mice, we found an increased expression of genes typically associated with immunosuppressive MAM (Arginase, CSF1R, CD206, Nos1, Ros1) (Fig. 2i, h), possibly derived from differentiation of initially cisplatin-recruited IM [33].

Pre-treatment of tumor-free SCID mice with cisplatin and a neutralizing antibody (nAb) against CCL2 prevented the BM expansion and lung recruitment of CCR2+CXCR4+ IM (Fig. 3a,b). Tail vain injection of H460 cells 72h post treatments revealed that anti-CCL2 nAb significantly reduced cisplatin-induced metastasis, MICs expansion and decreased the number of MAM (Fig. 3c-f), pointing at CCR2+CXCR4+ IM as the principal mediators of cisplatin pro-metastatic effects.

**Figure 3.**
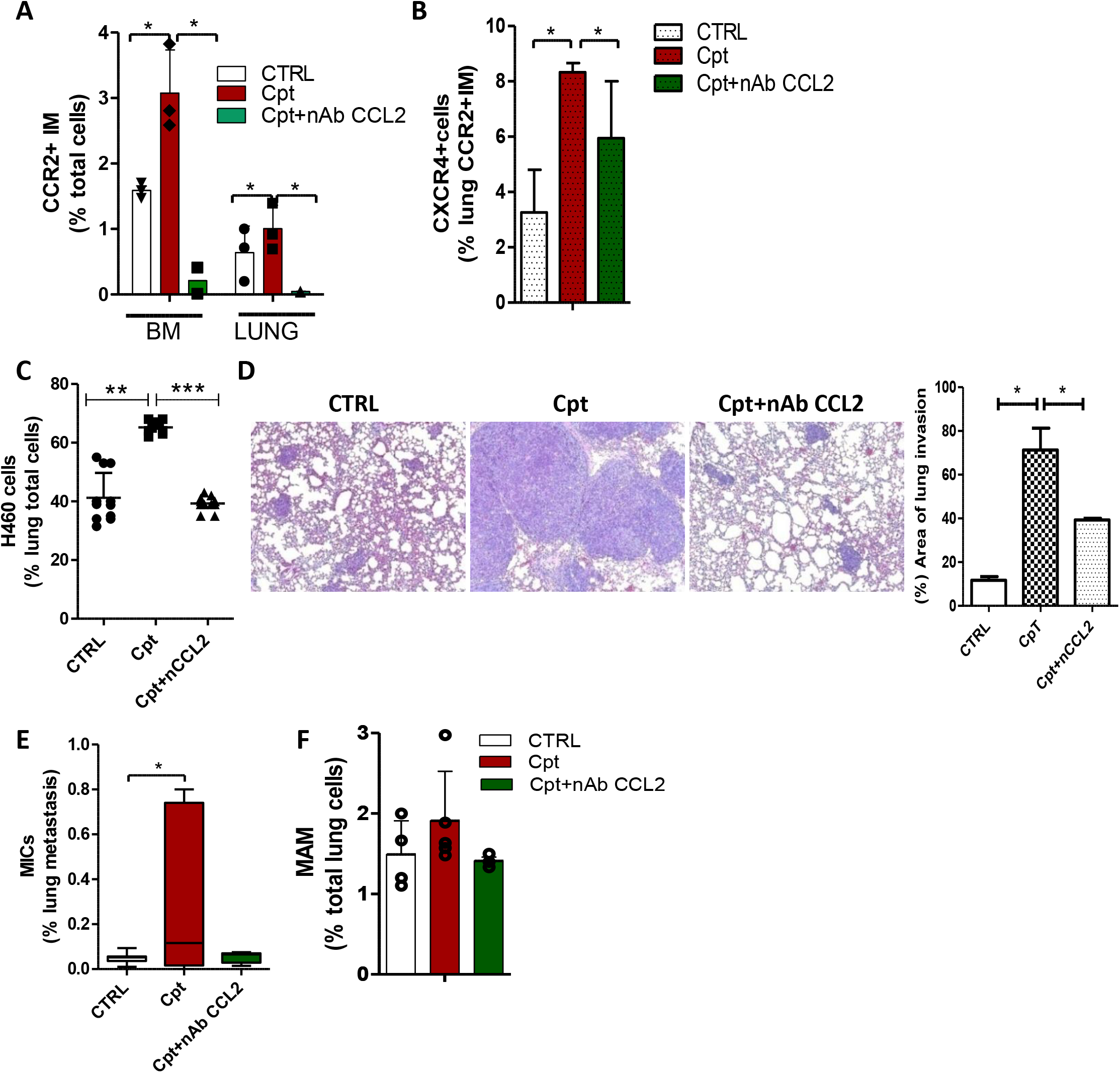
CCL2 inhibition counteracts cisplatin pro-metastatic effects. **A)** Percentage of CCR2+ inflammatory monocytes detected by FACS in SCID mice BM and lungs, 72h post treatment with cisplatin alone or with anti-CCL2 neutralizing antibody (nAb CCL2, 150μg/mice). N=3 mice/group * p≤ 0.05. **B)** Median frequency of CXCR4+ cells detected in lung CCR2+IM, analysed in A. N=3 mice/group * p≤ 0.05. **C)** Median percentage of lung metastatic H460 cells detected by FACS 3 weeks after i.v. injection in SCID mice pre-treated with cisplatin or combination with nAb CCL2. N= 6 lungs/group. ** p≤ 0.001. ***≤ 0.0001. **D)** H&E of H460 lung metastases in treatment groups described in (C) and histological quantification of the area of lung invasion by tumor cells (on the right N=6 lungs/group). *p≤ 0.05. **E)** Median percentage of CD133+ CXCR4+ MICs detected by FACS in H460 lung metastasis, analysed in (C). N= 6 lung/group. * p≤ 0.05. **F)** Percentage of metastasis-associated macrophages detected in metastatic murine lungs, analysed in C-D.

### Inflammatory monocytes increase tumor cells extravasation and expansion of MICs through SDF-1/CXCR4 axis

Chemotherapy can increase vascular permeability [16], that might be implicated in the early retention of tumor cells at the lungs, observed in cisplatin treated mice. Indeed, cisplatin treatment up-regulated in lung tissue several adhesion molecules able to mediate vascular adhesion and extravasation of leukocytes and tumor cells (ICAM-1, VCAM, Selectin-P) (Fig. 4a), data also confirmed *in vitro* in murine and human endothelial cells treated with cisplatin (Appendix Fig. S3a). This modulation was associated to the deconstruction of alpha-SMA+ endothelial layers, suggestive of an increased vascular leakiness (Appendix Fig. S3b).

**Figure 4.**
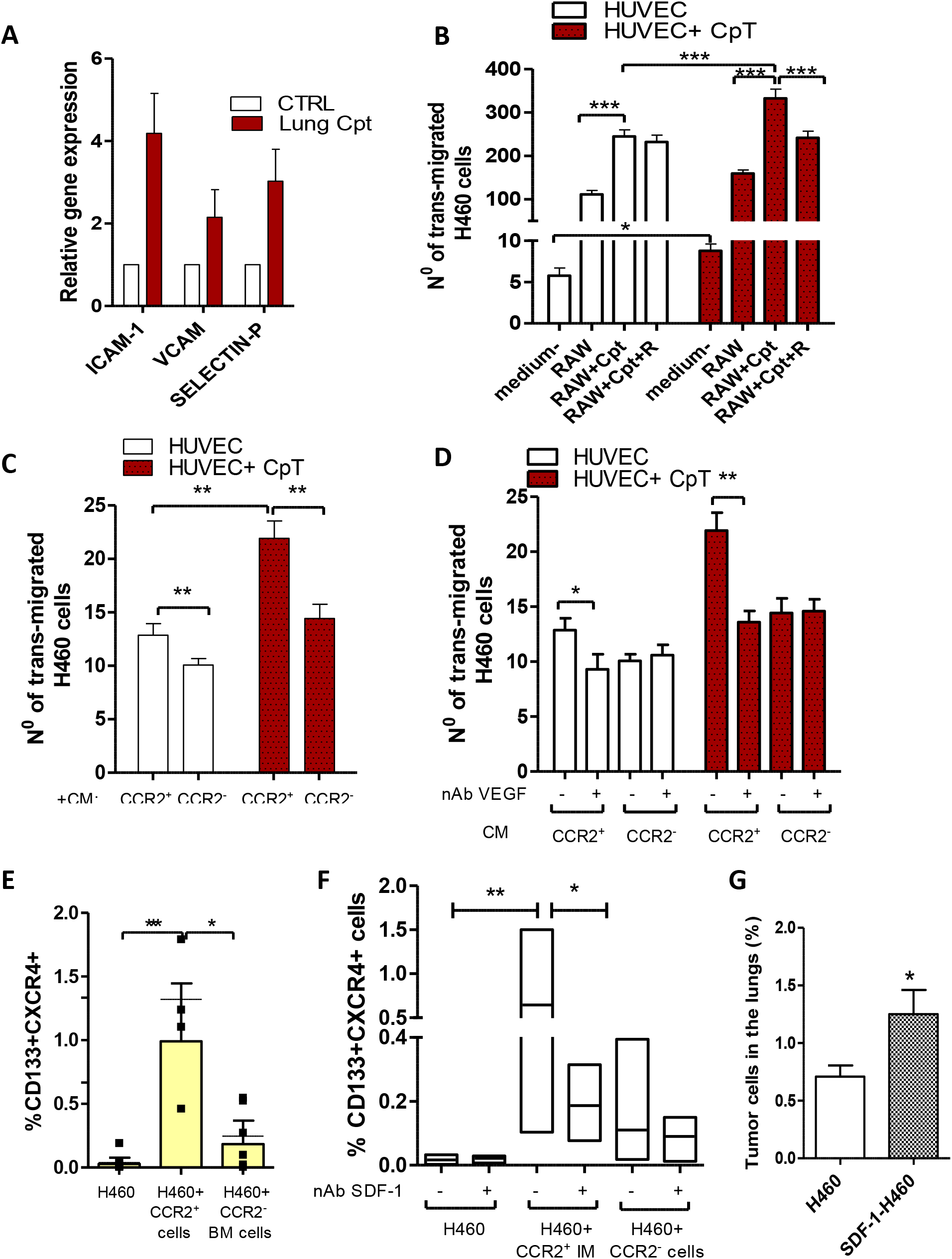
IM promote tumor cells extravasation and MICs expansion. **A)** Gene expression analysis of adhesion molecules performed n=4 SCID mice lungs, 72h post treatment with cisplatin. Untreated lung tissue was used as calibrator. **B)** Transendothelial migration assay: murine RAW 264.7 macrophages, treated with cisplatin (5μM) or combination with anti-CXCR4 (10μM), were used to chemoattract PKH labelled H460 cells trough endothelial cell layer, treated or not with cisplatin (5μM). Serum free medium was used to assess basal transmigration ability of tumor cells. Migrated PKH labeled H460 cells were counted by fluorescent microscope in 4 random fields of each insert, in duplicate. Bars are the mean ± SD of three independent experiments. * p≤ 0.05, ***≤ 0.0001. **C)** H460 cells were chemoattracted by conditioned medium (CM) from sorted bone marrow derived CCR2+ or CCR2-cells trough endothelial cell layer, treated or not with cisplatin, and analyzed as in (B). Bars are the mean ± SD of n=2 independent experiments. ** p≤ 0.001. **D)** Transendothelial migration assay as in (C), using CM from BM CCR2+ and – cells with or without neutralizing antibody (nAb) against VEGF (150ng/ml). Bars are the mean ± SD of n=2 independent experiments. * p≤ 0.05; ** p≤ 0.001. **E)** Percentage of CD133+CXCR4+ MICs evaluated by FACS in H460 cells co-cultured for 48h with BM-sorted CCR2+ and CCR2-cells. Bars are the mean ± SD of n=5 independent experiments. * p≤ 0.05; ** p≤ 0.001. **F)** Percentage of CD133+CXCR4+ MICs evaluated by FACS in H460 cells co-cultured with BM sorted CCR2+ and CCR2-cells with/ without nAb SDF-1 (25μM). Bars are the mean ± SD of n=3 independent experiments, each in technical duplicate. * p≤ 0.05; ** p≤ 0.001. **G)** Mean percentage of lung DTCs from subcutaneous xenografts ensuing from parental H460 and SDF-1 selected H460 cell lines. N= 4 mice/ group. * p≤ 0.05

In vitro trans-endothelial migration of H460 cells demonstrated that cisplatin-treated HUVEC endothelial cells allowed an enhanced transmigration of tumor cells chemoattracted by the murine macrophage cell line (RAW 264.7) (Fig. 4b). Moreover, RAW 264.7 macrophages treated with cisplatin showed a 10-fold enrichment in CCR2+CXCR4+ cell subset (Appendix Fig. S3c), that was associated with increased H460 transmigration. This phenomenon was inhibited by peptide R, able to prevent CCR2+CXCR4+ subset expansion (Fig. 4b).

We also confirmed that conditioned medium (CM) from BM-sorted CCR2+ IM increased the transmigration ability of H460 cells, especially through cisplatin-treated endothelial cells, compared to CM of CCR2-cells (comprised of neutrophils, NKs, resident monocytes and dendritic cells) (Fig. 4c).

We verified that compared to CCR2-myeloid cells, CCR2+IM expressed higher levels of VEGF (2.7 foldchange ± SD 1.2), known to mediate tumor cells extravasation at metastatic sites [13,14]. Coherently, cisplatin treated RAW 264.7 and murine lung tissues, both enriched in CCR2+CXCR4+ cells, up-regulated VEGF expression and neutralization of murine VEGF-A in CM of CCR2+ cells prevented tumor cell extravasation (Appendix Fig. S3d-e and Fig. 4d).

Next, we investigated the link between lung IM increase and MIC expansion caused by cisplatin treatment. Co-cultures of H460 cells with BM-sorted CCR2+ IM determined a 20-fold increase of CD133+CXCR4+ MICs compared to co-cultures with CCR2-myeloid cells (Fig. 4e). Interestingly, we verified that SDF-1 expression was 12.7-fold (±SD 7.5) increased in CCR2+ IM compared to CCR2-BM cells. Neutralization of SDF-1 specifically prevented the CCR2+ IM-induced MICs increase in H460 cells and also in lung adenocarcinoma cell lines (A549 and LT73), proving the ability of CCR2+ IM to foster MICs expansion via SDF-1/CXCR4 activation (Fig. 4f and Appendix Fig. S3f). Similarly, cisplatin-treated HUVEC endothelial cells, that also up-regulated SDF-1, promoted the expansion of CD133+ cells even if at a lesser extent than CCR2+ IM (Appendix Fig. S3g).

Finally, the key role of SDF-1 in mediating CD133+CXCR4+ expansion was functionally demonstrated by the high content of MICs (4.3 fold-increase) and enhanced ability to colonize murine lungs of an H460 subline selected by chronic exposure to SDF-1 (Fig. 4g).

### Cisplatin treatment of lung cancer xenografts expands MICs and recruits CXCR4+ tumor-associated macrophages (TAM) through SDF-1/CXCR4 axis promoting MICs intravasation

Treatments of H460 subcutaneous xenografts demonstrated that combination with peptide R did not enhance cisplatin efficacy in controlling tumor growth, as expected by the low percentage of CXCR4+ tumor cells within the tumor bulk (Appendix Fig. S4a). However, CXCR4 inhibitor prevented the enrichment in MICs content and the increase of tumor SDF-1 caused by cisplatin (Fig. 5a, b).

**Figure 5.**
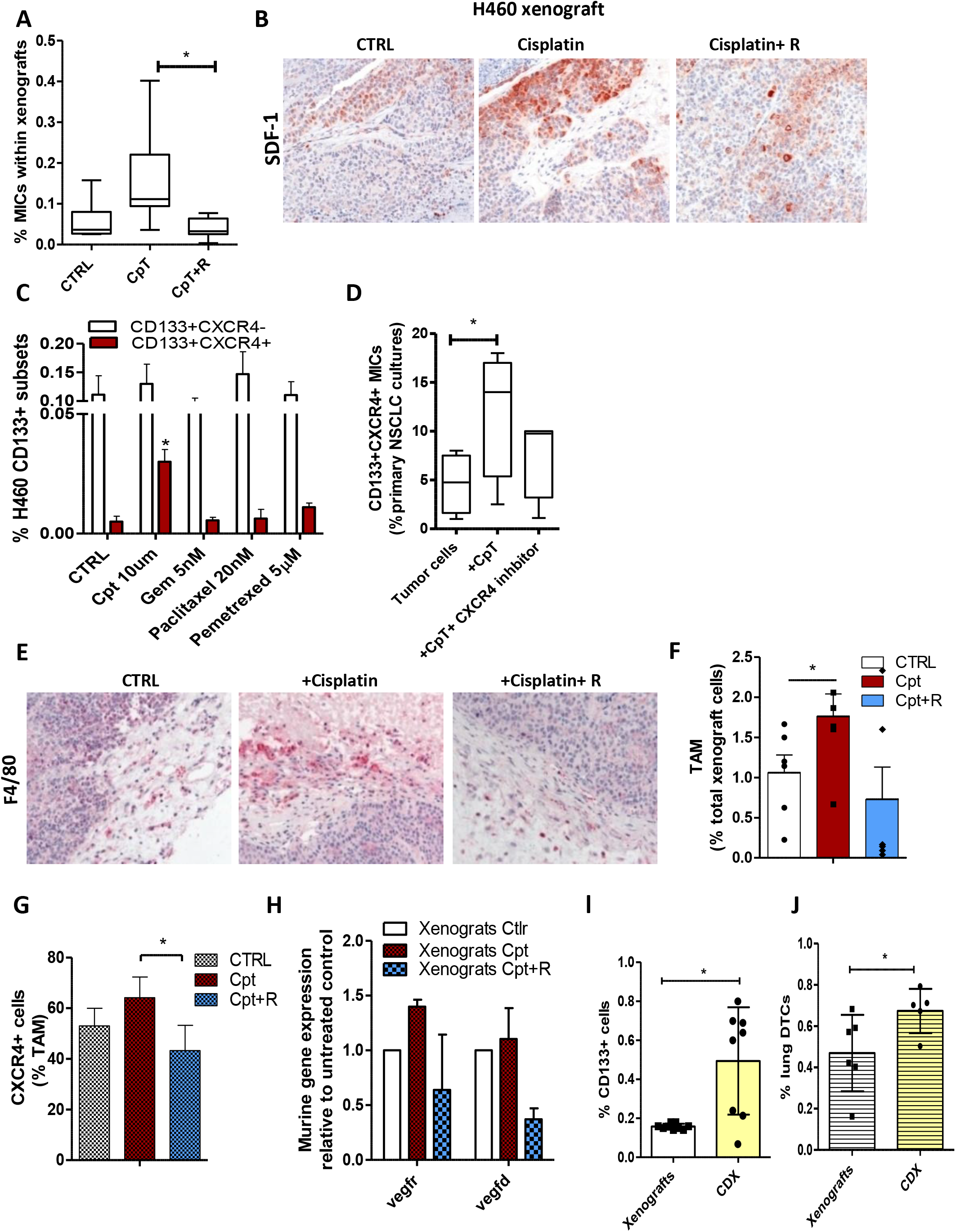
Cisplatin-induced SDF-1 in tumor cells favors MIC expansion and intravasation. **A)** Content of CD133+CXCR4+ MICs in H460 xenografts at the end of treatments. N= 4 mice/group, two independent experiments. * p≤ 0.05. **B)** IHC for SDF-1 in control and treated xenografts at the end of treatments. **C)** FACS analysis for CD133+ subsets evaluated in H460 cell line 72h after treatments with different chemotherapeutic drugs compared to untreated control. Bars are the mean ± SD of n=3 independent experiments. * p≤ 0.05. **D)** Median percentage of CD133+CXCR4 MICs evaluated by FACS in primary NSCLC cultures treated with cisplatin (10μM) and combination with CXCR4 inhibitor for 72h. N=4 independent experiments. *=p≤ 0.05. **E)** IHC for F4/80 macrophage marker in control or treated xenografts. **F)** Percentage of tumor associated macrophages (TAM) detected in xenografts at the end of treatment. Bars are the mean value ± SD. N= 3 mice/group, two independent experiments.* p≤ 0.05. **G)** Frequency of CXCR4+ cells within gated TAM, detected in (F). **H)** Gene expression analysis for murine *vegfr* and *vegfd* in n=3 treated xenografts (the same evaluated in F-G). Untreated xenografts were used as calibrator. **I)** FACS analysis for CD133+ cells evaluated in H460 xenografts and corresponding CTC-derived xenografts (CDX). Bars are the mean value ± SD of N=4 independent analysis, in duplicate.*p≤ 0.05. **J)** Percentage of disseminated tumor cells detected by FACS in lungs of mice bearing H460 xenograft and CDX evaluated in (I). Bars are the mean percentage ± SD. N=3 independent analysis, in technical duplicate.

Interestingly, *in vitro* treatment of H460 cells and others NSCLC cell lines (H1299/A549/LT73) with different chemotherapeutic drugs, at IC80 dose, demonstrated the unique ability of cisplatin to induce MICs expansion along with SDF-1 increase. Nevertheless, other chemotherapeutic drugs stimulated the production of different inflammatory cytokines able to expanded CD133+CXCR4-subset without modifying MICs (Fig. 5c and Appendix Fig. S4b-d). Finally, cisplatin treatment of short-term cultures of primary NSCLC confirmed MICs expansion that can be impaired by CXCR4 blockade (Fig. 5d).

Increased SDF-1 in cisplatin-treated xenografts also promoted the recruitment and the peritumoral accumulation of a specific subset of CXCR4+ tumor associated macrophages (TAM: F480^high^GR-1-CD206+), known to promote intravasation and angiogenesis via VEGF/VEGFR axis [17,19], that was prevented by peptide R (Fig. 5e-g). VEGF/VEGFR modulation was observed in treated xenografts, coherently with different content of CXCR4+ TAM (Fig. 5h).

We functionally proved that only circulating tumor cells (CTCs) isolated from blood of cisplatin-treated mice were able to generate tumors when injected subcutaneously in SCID mice. Ensuing CTC-derived xenografts (CDX) showed an increased content of CD133+ CSC compared to original xenografts and demonstrated an enhanced lung dissemination ability, likely reflecting the properties of circulating MICs originally expanded by cisplatin and allowed to escape primary tumors possibly through the interaction with CXCR4+TAM (Fig. 5i-j).

### Cisplatin treatment of lung cancer xenografts co-recruits MICs and metastasis associated macrophages (MAM) at distant sites through SDF-1/CXCR4 activation favoring metastasis formation

Cisplatin caused a boost in lung micro-metastases formation in mice bearing subcutaneous H460 xenografts, coupled with a significant enrichment in MICs content (1.8-fold change) (Fig. 6,a-c). We confirmed in cisplatin-treated mice an increased stromal SDF-1 expression in the lung parenchyma interstitium and a significant higher level of plasmatic CCL2 compared to control (Fig. 6d-e). Consistently, we observed an augmented lung recruitment of CCR2+CXCR4+ monocytes/macrophages (Fig. 6f, g), that can promote the extravasation and expansion of MICs subset, as we demonstrated above. Combination with CXCR4 inhibitor impaired SDF-1 increase, preventing the recruitment of CXCR4+ MICs and CXCR4+ MAM, overall resulting in diminished metastasis formation (Fig. 6a-g).

**Figure 6.**
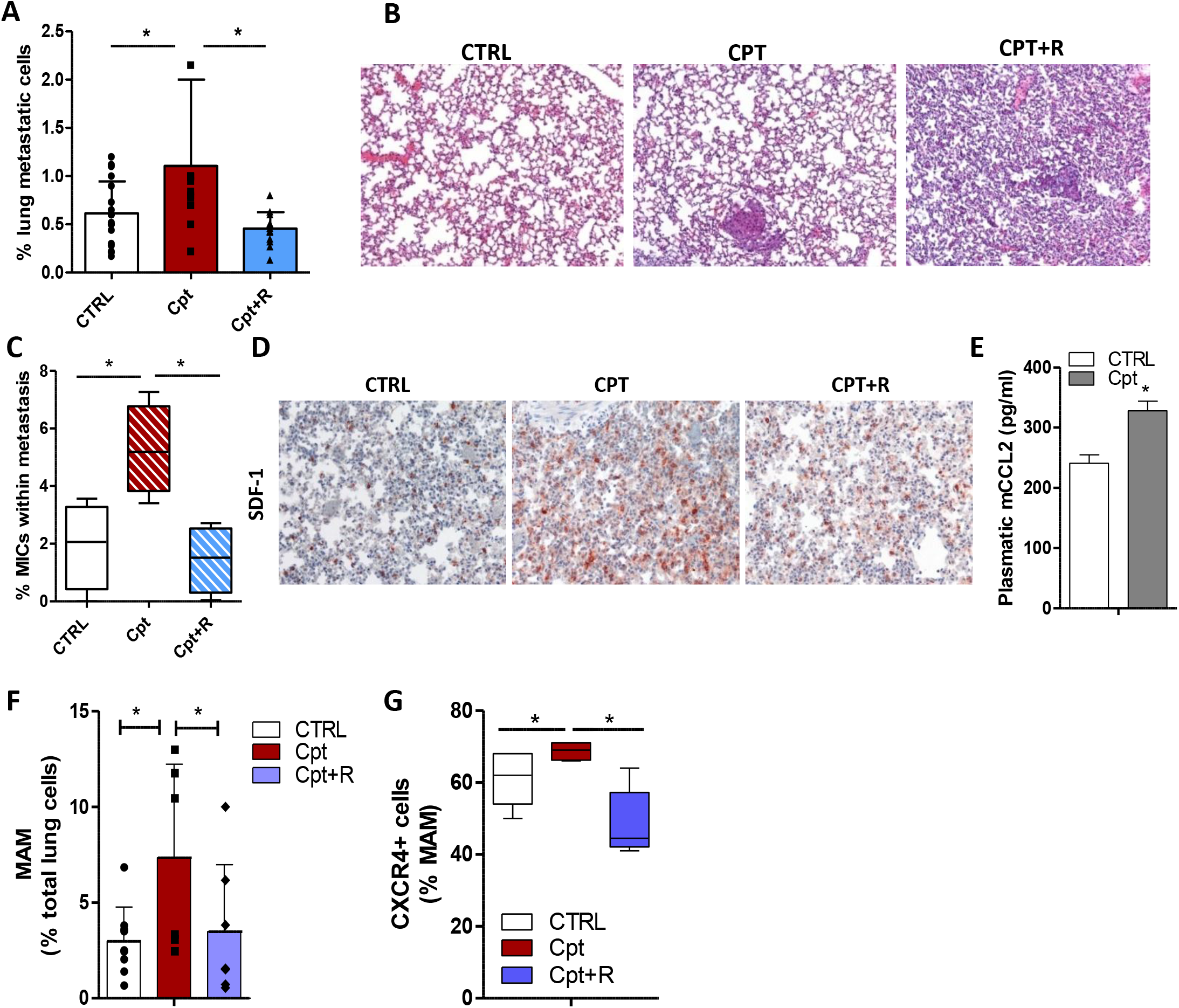
Cisplatin treatment fosters spontaneous metastasis formation co-recruiting MICs and IM. A) Percentage ± SD of H460 lung metastatic cells detected in mice-bearing subcutaneous xenografts treated with cisplatin or combination treatment (Fig. 6). N= 4 mice/group, two independent experiments. * p≤ 0.05. B) H&E of lungs of mice-bearing H460 xenografts analysed in (A). C) CD133+CXCR4+ MICs content analysed by FACS in H460 lung metastasis (as detected in A). N=4 mice/group, two independent experiments * p≤ 0.05. D) IHC for SDF-1 in lungs of control and treated mice bearing H460 xenograft. E) ELISA quantification of CCL2 in plasma of mice-bearing xenografts, at the end of treatments. Bars are the mean value ± SD of duplicate experiments, performed on plasma sample, pooled from N=2mice/ group. F) Percentage of metastasis associated macrophages (MAM) detected in murine lungs analysed in C. Bars are the mean value ± SD. N= 4 mice/group in independent duplicate experiments.* p≤ 0.05. G) Frequency of CXCR4+ MAM detected in (F) * p≤ 0.05

### SDF-1 levels are increased in platinum-treated clinical samples and correlate with poor clinical outcome

To evaluate the clinical relevance of our pre-clinical data, we assessed by immunohistochemical staining the expression of SDF-1 in surgical specimens from chemo-naive NSCLC patients (n=57) and NSCLC patients treated with platinum-based chemotherapy in the neoadjuvant setting (n=57), matched for stage (IIIA), histological subtypes, sex and age (Appendix Table S1). The frequency of SDF-1 positive tumors was significantly higher in the neoadjuvant-treated group than in untreated tumors (67% vs. 26%; p>0,0001).

To address a possible prognostic role for SDF-1, patients treated with neoadjuvant platinum-based chemotherapy (n=39, Appendix Table S2) were categorized according to the SDF-1 IHC score (percentage of positive cells X staining intensity) (Fig. 7a): patients with tumor SDF-1 score>6 exhibited a significant shorter disease-free survival (DFS) (p=0,0056; Hazard Ratio= 3,1) and overall survival (OS) (p=0,029; Hazard Ratio= 3,46) compared to patients with SDF-1 score<6 (Fig. 7b).

**Figure 7.**
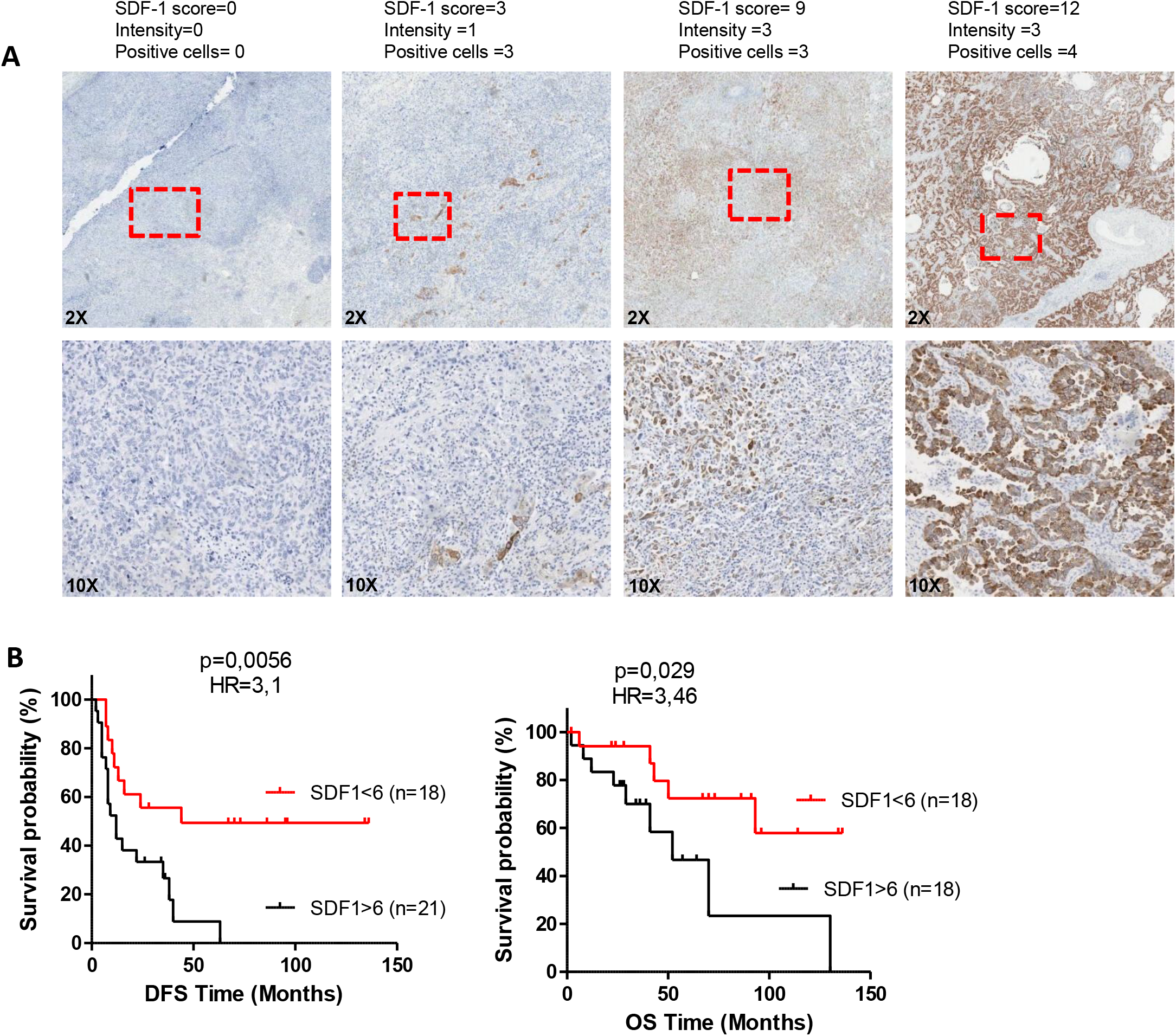
High SDF-1 levels correlate with poor prognosis in NSCLC patients treated with neoadjuvant cisplatin. **A)** IHC for SDF-1 of platinum-based neo-adjuvant treated NSCLC tumors, representative of different score calculated as: %positive cells x Intensity. Areas within dashed lines are shown at higher magnification. **B)** Kaplan-Meier curves for DFS and OS respectively of n=39 and n=36 NSCLC patients treated with platinum-based neo-adjuvant therapy, categorized according to SDF-1 tumor score>6. Reported *p* values are calculated by Log-rank (Mantel-Cox) Test.

## Discussion

Herein we demonstrate the crucial role of CXCR4/SDF-1 axis activation both in stromal and tumor cells in mediating the paradoxical pro-metastatic activity of cisplatin. Early responses to cisplatin insults include the expansion in the bone marrow of CCR2+CXCR4+ inflammatory monocytes (IM) and their recruitment to the lungs (the most common metastatic site for NSCLC)[34], guided by the increased stromal release of SDF-1 and CCL2, that can be efficiently prevented by CXCR4 blockade using a novel peptide inhibitor (peptide R) [29].

The CCR2+ subset of IM has already been described as an important player in early phase of metastasis formation and tumor cells survival [14,32,35] and also may function as monocytic-myeloid derived suppressor cells (M-MDSC) in mice [31]. The subset of CCR2+ IM can favor the extravasation of tumor cells at distant sites through the release of vascular endothelial growth factor (VEGF-A) (23) and supporting their growth [36]. Notably, chemotherapy may exacerbate tumor release of CCL2 and consequently the recruitment of CCR2+IM to both primary and metastatic sites [37]. Interestingly, extracellular vesicles released by chemotherapy-treated murine breast cancer cells promoted endothelial cell release of CCL2 and consequent recruitment of CCR2+ IM at the lung pre-metastatic niche, overall fostering metastasis development [15].

Our data corroborate the pro-metastatic effects of CCR2+ IM and in addition show that, compared to other chemotherapeutic drugs used for NSCLC treatment, cisplatin uniquely chemo-attracts to the pre-metastatic niche the population of CCR2+ IM co-expressing CXCR4. This observation was confirmed both in SCID and immunocompetent mice, proving the central role of the innate immune responses in mediating cisplatin pro-metastatic effects, as also previously reported [38]. Moreover, our data support the pro-tumorigenic/metastatic properties of CXCR4 myeloid cells, already verified in a transgenic mouse model with genetic deletion of CXCR4 in the myeloid compartment [39].

Our data in tumor-free mice, also indicate that acute response to cisplatin may preferentially trigger the generation inflammatory status and expansion of IM rather than M-MDSC, that share similar phenotype but possess different immunosuppressive properties [40]. Then, the interactions with metastatic cells can induce the differentiation of IM into immunosuppressive MAM, as already described in literature [33].

We show that both CCL2 and CXCR4 inhibition in combination with cisplatin can impair IM recruitment and prevent cisplatin-induced metastasis formation. Although CCL2 blocking impairs metastasis formation [13,14,32], cessation of CCL2 neutralization *in vivo* can cause a massive release of BM monocytes that, paradoxically, exacerbate metastasis development by promoting cancer cell mobilization and blood vessel formation [13]. Altogether, these observations point at CXCR4 inhibition as a potentially safer and effective strategy to be exploited, in combination with cisplatin, to prevent pro-metastatic CXCR4+ IM mobilization.

We verified, both in vivo and in *vitro,* that cisplatin can induce endothelial leakiness favoring tumor cells extravasation, data in line with reported altered vascular permeability and overexpression of VEGFR-1 caused by chemotherapy, associated with augmented lung metastasis [16,37,38]. Moreover we showed that the increase of IM recruited by cisplatin can additionally foster tumor cells trans-endothelial migration via VEGF release, in accordance to previously reported observations [14].

To the best of our knowledge this is the first time that IM, recruited by cisplatin to the lungs, are reported to select/expand MICs by releasing high levels of SDF-1 overall fostering metastasis development.

At primary tumor site, cisplatin specifically selects chemoresistant MICs subset [26,27]: cisplatin-induced tumor SDF-1 increase can contribute to expand MICs through the activation of CXCR4 pathway and also can possibly convert non-MICs into MICs, as we recently demonstrated (6).

Among different chemotherapeutic drugs routinely used for NSCLC treatments, we show here that only cisplatin induces SDF-1 expression and consequently can expand CXCR4+ MICs. Nevertheless, other chemotherapeutic agents can expand the fraction of CD133+CXCR4-CSC, in line with previous data [41], by inducing the release of different inflammatory and pro-tumorigenic cytokines, as described in several reports [7,10,16,42].

Cisplatin-induced increase of SDF-1 also fosters the recruitment at primary tumor site of CXCR4+ tumor associated macrophages (TAM) that have been shown to favor tumor cells intravasation, neo-angiogenesis, lymphangiogenesis and to promote tumor relapse after chemotherapy [19,43–45].

Notably, we provide functional proof-of-concept that the expansion of MICs at primary site along with CXCR4+ TAM increase enhances the ability of tumor cells to escape primary tumor, as assessed by the unique ability of CTC isolated from blood of cisplatin treated mice to generate tumors when transplanted in new recipient mice.

Finally, we prove in clinical samples from NSCLC patients that SDF-1 is significantly increased in tumor cells after platinum-based neo-adjuvant chemotherapy and, remarkably, that a high SDF-1 expression in treated patients is correlated with shorter progression free and overall survival.

Very few studies have investigated the prognostic value of SDF-1 in primary NSCLC and among them no significant association between SDF-1 expression and DFS or OS was reported [46,47]. Our data prove the prognostic relevance of SDF-1 marker in the setting of neo-adjuvant treated NSCLC patients, even if a prospective longitudinal study is needed to confirm the clinical value of our preliminary observations.

Taken together our findings prove that cisplatin uniquely activates the SDF-1/CXCR4 axis, in both stromal and tumor cells, resulting in multiple metastasis-promoting actions. A novel use of CXCR4 blockade in combination with cisplatin may control more effectively the metastatic disease in NSCLC patients and, moreover, could be exploited in chemotherapy/ICI doublets regimen, the future mainstay of advanced NSCLC therapy (3, 32, 46), to extend the benefit of immunotherapy potentially turning to advantage the immunomodulatory effects of cisplatin.

## Materials and Methods

### Cell cultures and reagents

H460, A549, H1299 tumor cell lines, authenticated by STR profiling, murine RAW 264.7, murine 3B11 and human HUVECs endothelial cells (all purchased from ATCC) were cultured in RPMI 1640+ 10% Fetal Bovine Serum (FBS) or in endothelial grown media (EGM-2-Lonza).

CCR2+ cells were positively isolated from BM cells flushed from SCID mouse femurs using anti-CCR2-APC+ Anti-APC MicroBeads and autoMACS^®^ Pro Separator (Miltenyi). 1×10^5^ sorted CCR2+ IM or CCR2-myeloid cells were co-cultured at ratio 1:1 with tumor cells or used to recover conditioned medium after 48h.

Short-term NSCLC primary cultures were obtained by differential filtration of dissociated primary tumor tissue, as described in Bertolini *et al* [26], from a consecutive series of Caucasian male and female of early stage (I-IIb) NSCLC patients undergoing surgical resection.

For in vitro experiments, cells were treated with: peptide R 10μM; recombinant human SDF-1α 25 ng/ml (300-28A), CytoMCP-1 (CCL2) 25ng/ml (300-04), IL-8 50ng/ml (200-08), GRO-α/MGSA 25ng/ml (300-11), IL-6 10ng/ml (200-06) (all from Peprotech); neutralizing Ab anti-human/mouse SDF-1 25μg/ml (MAB310) and mouse VEGF 150ng/ml (VEGF164) (all from R&D System).

### Flow cytometry

Immune cell subsets (live cells/CD45+) were identified in BM and lungs as: low SSC/DX5+ NKs; CD11b+/LY6G+ neutrophils; Ly6G-/Ly6C^-/low^/F480^high^/CD11b-/CD11c+ alveolar macrophages; Ly6G-/Ly6C^-/low^/F480^high^/CD11b+CD11c-monocyte-derived macrophages;Ly6G-/Ly6C-/F480^-/low^/CD11b+/CD11c+ conventional dendritic cells; Ly6G-/F480^-/low^/CD11b+/CD11c-/Ly6C^High^ inflammatory monocytes or Ly6C^dim^ resident monocytes. Expression of CXCR4 and CCR2 was evaluated within mono/macrophages subsets.

Lung disseminated/metastatic tumor cells were indentified in lung dissociated tissues by FACS as 7-AADneg/mouse MHC class I neg cells [26]. List of all antibodies is available in Appendix Supplementary Methods. Gallios (Beckman Coulter) or FACSCanto (BD Bioscience) flow cytometers were used for data acquisition and FlowJo software V10 (RRID:SCR_002798) for data analysis.

### Trans-endothelial migration assay

5×10^4^ HUVEC cells were plated in EGM-2 medium into the upper chamber of FluoroBlok 24 well cell culture inserts (Corning) covered with 100μl of Matrigel (Corning). In the lower chamber 5×10^4^ RAW 264.7 were plated in RPMI+10% FBS or filled with conditioned medium from CCR2+ and CCR2-BM cells. The following day HUVEC and RAW cells were treated with cisplatin 5μM plus/minus peptide R 10μM. 2×10^4^ PKH-26 (Sigma) stained H460 cells were plated onto the HUVEC endothelial cells layer and after 48h migrated H460 cells were fixed with PFA 4% and counted under a fluorescence microscope.

### Animal studies

For lung conditioning experiments, naïve female SCID mice and Balb/c mice (Charles River Laboratories) were treated with i.p. cisplatin (Teva Pharma) 2,5mg/kg followed by i.p. administration of peptide R (2mg/kg), once every 3 days and next injected i.v. with 1×10^6^ H460 cells. Treatment schedules for other chemotherapeutic drugs are available in Appendix Supplementary Methods

Anti-mouse CCL2 antibody (150μg/mice) (clone: 2H5 BioXCell) was administered to SCID mice together with cisplatin and for the following 3 days, before H460 cells injection.

Mice with palpable xenografts (TW>=50mm^3^), generated by s.c. injection of 1×10^5^ cells H460 cells with Matrigel (Corning) were randomized into 3 groups: untreated control, treated with 5mg/kg cisplatin (i.p) every 7 days alone or with 2mg/kg peptide R (i.p.) once every 5 days post cisplatin for 3 weeks. *In vivo* experiments protocols were revised by the Internal Ethics Committee for Animal Experimentation and approved by the Italian Ministry of Health.

### SDF-1 IHC score

NSCLC specimens were obtained from consenting patients (clinicopathological characteristics are available in Appendix Table S1-2) undergoing surgical tumor resection after platinum-based neo-adjuvant chemotherapy (study INT 201/18 approved by the Independent Ethics Committee). IHC score for tumor SDF-1 staining (Clone 79018 R&D Systems, IHC staining details are available in Appendix Supplementary Methods) was obtained by multiplying the percentage of positive cells (P=0 no positive cells; P=1 1-25% positive cells; P=2 25% −50% positive cells; P=3 50-75% positive cells; P=4 > 75% positive cells) by the staining intensity (I=1-3). The pathologist M.M. performed the histological blinded evaluation of SDF-1 staining.

### Statistical Analyses

Statistical analyses were performed using GraphPad Prism v5.0 (RRID:SCR_002798). Statistically significant differences were determined with Student t tests when comparing two groups or ANOVA test for multiple comparisons. Sample size and number of replicates were indicated in Figure legends for each experiments. Data are presented as mean (±SD), unless otherwise indicated. SDF-1 expression association with categorical clinical-pathological characteristics of NSCLC patients was calculated with Pearson’s chi-squared test. Survival data were analyzed using Kaplan-Meier log-rank tests. All tests were two-sided and the statistical significance was defined as a *P* value less than .05.

## Availability of data and materials

All data generated or analysed during this study are included in this published article and its supplementary information files

## Authors’ contributions

GB, LR, GS, SS, CT, NZ conceived the study. GB, VC, OF, MT, GC, CC, FF, FG, MM, CC, CDA performed the experiments and analyzed the data. MM performed pathological evaluation of human specimens. CT performed pathological evaluation of murine specimens. GLR, ADT, UP selected case series and collected clinical information. GB, LR, SS wrote the manuscript.

